# *Klebsiella pneumoniae* exhibiting a phenotypic hyper-splitting phenomenon including the formation of small colony variants

**DOI:** 10.1101/2024.01.11.575232

**Authors:** Eyüp Doğan, Katharina Sydow, Stefan E. Heiden, Elias Eger, Georgi Wassilew, Richard A. Proctor, Jürgen A. Bohnert, Evgeny A. Idelevich, Katharina Schaufler, Karsten Becker

**Author notes:** shared last (Katharina Schaufler, Karsten Becker).

## Abstract

In this study, we characterized a *Klebsiella pneumoniae* strain in a patient with shrapnel hip injury, which resulted in multiple phenotypic changes, including the formation of a small colony variant (SCV) phenotype. Although already described since the 1960s, there is little knowledge about SCV phenotypes in *Enterobacteriaceae*. The formation of SCVs has been recognized as a bacterial strategy to evade host immune responses and compromise the efficacy of antimicrobial therapies, leading to persistent and recurrent courses of infections. In this case, 14 different, clonally identical resisto- and morpho-types were distinguished from the patient’s urine and tissue samples. Whole genome sequencing revealed the *K. pneumoniae* high-risk clonal lineage belonging to sequence type 147. Subculturing the SCV colonies consistently resulted in the reappearance of the initial SCV phenotype and three stable normal-sized phenotypes with distinct morphological characteristics. Additionally, an increase in resistance was observed over time in isolates that shared the same colony appearance. Our findings highlight the complexity of bacterial behavior by revealing a case of phenotypic “hyper-splitting” in a *K. pneumoniae* SCV and its potential clinical significance.

## Introduction

*Klebsiella pneumoniae*, an opportunistic pathogen known for its ability to cause a wide range of nosocomial and community-acquired infections, has emerged as a significant public health threat due to its strain-specific, extensive arsenal of resistance and virulence factors (1, 2). Infections caused by multi-, extensively-, and pandrug-resistant strains result in high mortality due to limited response to antibiotic therapy, which poses an increasing threat (3-5). Apart from classic strains, a hypervirulent *K. pneumoniae* (hvKp) pathotype occurs and is characterized by invasive, often life-threatening and multiple site infection, characteristically in healthy patients from the general population (6). In addition, convergent types that successfully combine resistance and hypervirulence represent a “perfect storm” and have been increasingly reported in recent years (7-9).

Beyond typical resistance mechanisms against various antimicrobials, functional resistance mechanisms have been elucidated that lead to antimicrobial treatment failure and foster the development of relapses and persistent infections (10). The formation of a biofilm matrix represents one of these mechanisms that facilitates antibiotic tolerance and the generation of bacterial persister cells (10). Interestingly, it has been demonstrated that a decrease in capsule biosynthesis, which is crucial for hypervirulent phenotypes, leads to increased *in vitro* biofilm formation and intracellular persistence (11). Another non-classical mechanism leading to functional resistance is the formation of the small colony variant (SCV) phenotype. SCVs are subpopulations of bacteria that exhibit slow growth, reduced colony size, and altered phenotypic properties compared to their normal-growing counterparts, making them difficult to detect and treat effectively (12, 13). Their ability to evade the host’s immune surveillance and to undermine the effectiveness of antimicrobial interventions by host cell internalization results in intracellular persistence, which contributes significantly to the recurrence and chronicity of the infection (14, 15). Another pivotal attribute facilitating this phenomenon is their capability to modulate metabolic processes and virulence characteristics (16, 17).

Hypermutator SCVs characterized by higher mutation frequencies than wild-type strains and isolated especially from cystic fibrosis (CF) patients (18, 19) have also been associated with antibiotic resistance (20, 21) and biofilm formation (22).

To date, research has focused on staphylococcal SCVs, while SCVs of Gram-negative bacteria have been investigated in only a few studies and case reports (12). Although the formation of small colonies in *K. pneumoniae* has been noticed during resistance studies against cephalosporins in the mid-1960s (23), this issue has not received sufficient attention and detailed research has not been conducted on this subject. The first clearly defined SCV of *K. pneumoniae* (SCV-Kp) in literature was obtained by *in vitro* exposure to gentamicin (24). SCV-Kp were also isolated from a patient treated with aminoglycoside antibiotics (25). Smaller and non-mucoid colonies were obtained as a result of conjugation-induced mutation in the outer membrane protein of a hypervirulent *K. pneumoniae* isolate (26). Another study showed that biofilm-forming *K. pneumoniae* developed heteroresistance to colistin by presenting slow-growing SCV-Kp (27).

Here, we report on *K. pneumoniae* isolates displaying 14 different resisto- and morpho-types obtained from an immunocompetent male patient, who had sustained a traumatic injury caused by shrapnel shell fragments. The isolates comprise an initial, mostly susceptible *K. pneumoniae* isolate with typical morphological characteristics isolated from the patient’s urinary specimen. From the urine and tissue samples, 13 additional phenotypes with different combinations of resistance and morphological characteristics including *K. pneumoniae* SCV phenotypes were isolated.

## Methods

### Patient data

Sufficient information could not be obtained regarding the period from the patient’s first acetabular and femoral head shrapnel-caused war injury in Ukraine in March 2022, where he underwent hip prosthesis at an external center before his transfer to our orthopedic service in July 2022. Fracture-related joint infection treatment in our hospital continued through November 2022. The administration of antibiotics during this period included piperacillin/tazobactam from July to October, 2022, trimethoprim/sulfamethoxazole from July to August, 2022, cefiderocol from August to November, 2022, and colistin from October to November, 2022. Daptomycin was introduced into the treatment protocol starting from October 2022 upon detection of *Staphylococcus epidermidis* from tissue samples and central venous catheter tip, and continued until the patient’s discharge. Subsequently, a planned course of post-discharge antibiotic suppression therapy with doxycycline for three months was initiated. The first identification of carbapenem-resistant *K. pneumoniae* (CRKP) occurred in July 2022, followed by the initial detection of SCV-Kp in September 2022. Therefore, we decided to aggregate and systematically assess the entirety of *K. pneumoniae* strains isolated from the patient.

### Strain identification

The urine sample obtained from the patient was quantitatively inoculated onto a Columbia agar plate with 5% sheep blood (BD Diagnostics, Heidelberg, Germany) and a MacConkey II-Agar plate (BD Diagnostics) using a 10 μl disposable sterile loop. The plates were then incubated for 48 hours. Tissue samples collected during surgery were inoculated onto Columbia agar plates with 5% sheep blood, MacConkey II-Agar plates, and Mueller Hinton Chocolate agar plates (all from BD Diagnostics). These plates were incubated under capnophilic conditions for up to seven days. The remaining tissue material was inoculated onto Schaedler agar and into BBL Fluid Thioglycollate media (both from BD Diagnostics) and incubated for up to 14 days under anaerobic and capnophilic conditions, respectively.

Preliminary characterization of each phenotype was grounded in colony morphology and minimal inhibitory concentration (MIC) results for antibiotics encompassed within the VITEK® 2 AST card specific to *Enterobacterales* (bioMérieux SA, Marcy l’Étoile, France) according to EUCAST criteria. All *K. pneumoniae* strains, isolated from various patient’s specimens during the period from July to December 2022, were identified by matrix-assisted laser desorption/ionization time-of-flight mass spectrometry (MALDI-TOF MS) utilizing the MALDI Biotyper® sirius system (Bruker Daltonics, Bremen, Germany) with MBT Biotargets 96 (Bruker Daltonics). The presence of carbapenemase-encoding genes was verified by a loop-mediated isothermal amplification (LAMP)-based assay (eazyplex®, AmplexDiagnostics, Gars-Bahnhof, Germany).

### Characterization of the phenotypes

Sequential subcultures of all phenotypic variants were carried out on various agar plates (including Columbia agar + 5% sheep blood, MacConkey agar from BD, and CHROMID® CPS® Elite agar from bioMérieux) to observe whether changes in colony morphology occurred and SCVs remained stable, followed by meticulous analysis of generated phenotypic profiles.

In order to determine colony sizes, each phenotype was inoculated onto 5% sheep blood agar plates in triplicate on different days. After overnight incubation at 35±1°C in ambient air, the diameters of ten colonies were measured on each plate and mean values were determined.

### Antimicrobial susceptibility testing

In addition to the initial VITEK^®^ 2 AST, the MICs of a standardized set of antibiotics were determined by the broth microdilution (BMD) method using cation-adjusted Mueller–Hinton broth (CAMHB; Micronaut-S 96-well microtiter plates, Merlin, Bornheim-Hersel, Germany), and for cefiderocol using iron-depleted CAMHB (UMIC®, Merlin, Bornheim-Hersel, Germany), as recommended by ISO 20776-1, the European Committee on Antimicrobial Susceptibility Testing (EUCAST), and the Clinical and Laboratory Standards Institute (CLSI) guidelines (28-30). The results were observed following 18±2 hours of incubation at 35±1°C in ambient air. All tests were conducted in triplicate on different days, and median MIC values were computed for analysis. *Escherichia coli* ATCC 25922, *E. coli* ATCC 35218, *K. pneumoniae* ATCC 700603, and *Pseudomonas aeruginosa* ATCC 27853 were used as quality control (QC) strains, and their results were within the QC range throughout the study. EUCAST Clinical Breakpoint Tables v. 13.1 were used for MIC interpretation (31).

### DNA isolation and sequencing

After overnight growth on blood agar plates at 37 °C, ten colonies were randomly selected and suspended in 1.5 mL tubes (Carl Roth, Karlsruhe, Germany) with 1 mL of phosphate buffered saline. Total DNA was extracted using the MasterPure DNA Purification kit for Blood, v. 2 (Lucigen, Middleton, WI, USA) according to the manufacturer’s instructions. Quantification of isolated DNA was performed with the Qubit 4 fluorometer and the dsDNA HS Assay kit (Thermo Fisher Scientific, Waltham, MA, USA). DNA was sent to SeqCenter (Pittsburgh, PA, USA), where sample library preparation using the Illumina DNA Prep kit and IDT 10bp UDI indices was performed. Subsequently, libraries were sequenced on an Illumina NextSeq 2000, producing 2×151bp reads. Demultiplexing, quality control and adapter trimming at the sequencing center was performed with bcl-convert v. 3.9.3 (https://support-docs.illumina.com/SW/BCL_Convert/Content/SW/FrontPages/BCL_Convert.htm).

### Assembly and genomic characterization

We employed a custom assembly and polishing pipeline to assemble raw sequencing reads to contigs. This pipeline consists of four parts, namely trimming (BBDuk from BBTools v. 38.98 [https://sourceforge.net/projects/bbmap/], quality control (FastQC v. 0.11.9 [https://www.bioinformatics.babraham.ac.uk/projects/fastqc/]), assembly (shovill v. 1.1.0 [https://github.com/tseemann/shovill]) with SPAdes v. 3.15.5 (32), and polishing (BWA-MEM2 v. 2.2.1 (33), Polypolish v. 0.5.0 (34)).

Genotyping was performed with Kleborate v. 2.2.0 (35) and Kaptive (36, 37).

### Confirmation of clonality

Trimmed sequencing reads of all isolates were mapped against isolate 1-A with snippy v. 4.6.0 (https://github.com/tseemann/snippy) and the SNP distance matrix calculated with snp-dists v. 0.8.2 (https://github.com/tseemann/snp-dists).

## Results

Overall, 14 distinct phenotypes were determined (Table 1). From the urine, two phenotypes (1-A and 1-B) exhibiting a normal colony size and glistening surface but differing in the color of their colonies displaying whitish or grey colonies, were isolated. All other phenotypes (n = 12) were isolated from tissue specimens. Strains numbered 1-A, 2-A, 3-A, 4-B, 5-B, numbered 1-B, 2-B, 3-B, 4-C, 5-C, and numbered 4-D, 5-D, displayed identical morphological attributes each, distinguished by whitish, glistening, and smooth (Figure 1-A), grey, glistening, and smooth (Figure 1-B), and grey, dry, and rough colonies (Figure 1-C), respectively. These strains revealed a normal colony size of 2.4 mm on average (range, 1 – 5.5 mm). The isolates displaying the SCV phenotype, numbered 4-A and 5-A, exhibited similar morphological characteristics, and colony sizes were smaller than 0.5 mm (Figure 1-D). No discernible variation in terms of colony clustering was observed among the various agar plates.

**Table 1.**
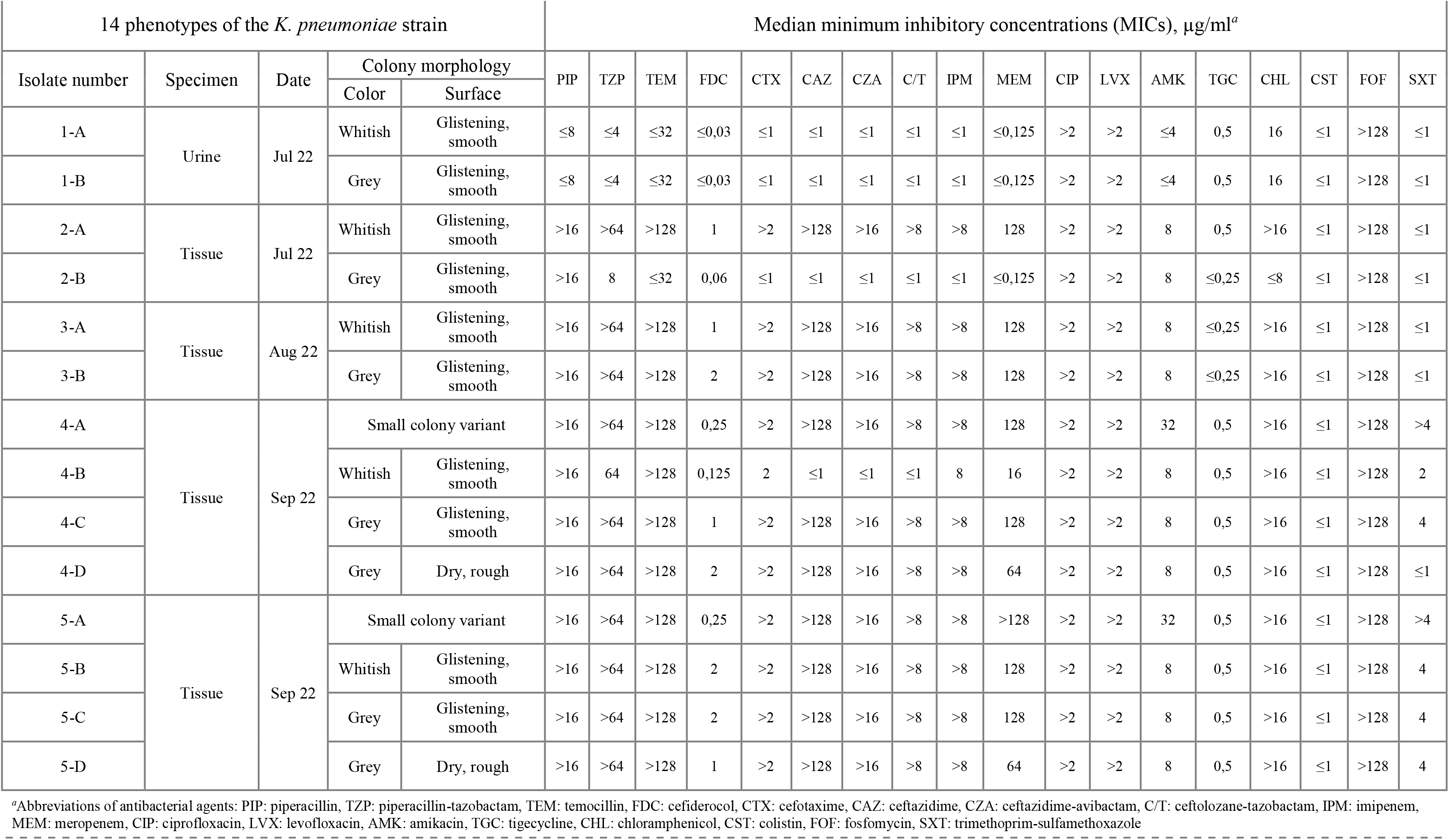
Colony morphology and antimicrobial susceptibility characteristics of the 14 phenotypes of the *Klebsiella pneumoniae* strain.

**FIG 1.**
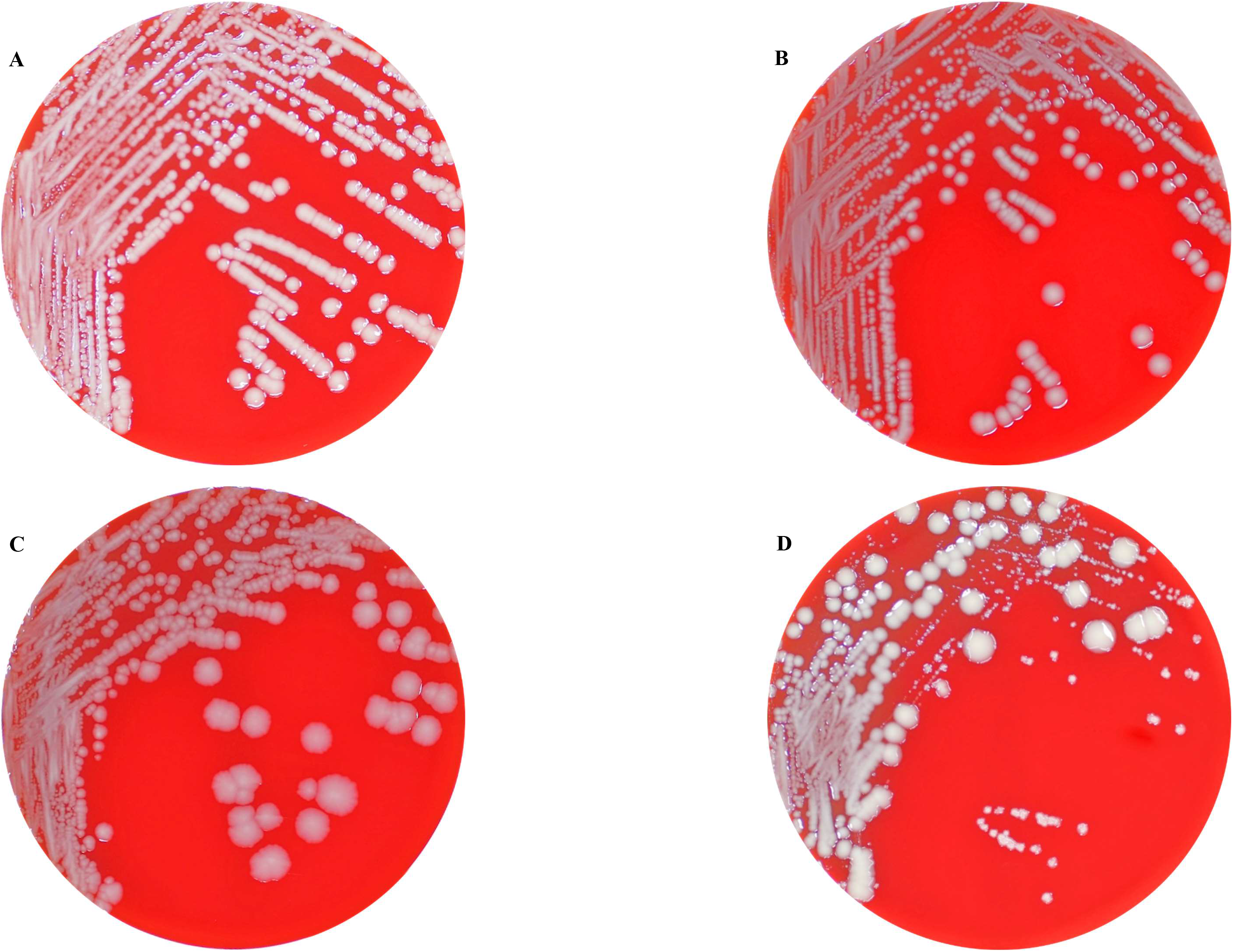
Columbia blood agar plates showing the different colonial morphotypes of the *K. pneumoniae* isolates comprising regular sized colonies (wild-type) with glistening whitish (Figure 1-A) and grey (Figure 1-B), and dry and rough grey colonies (Figure 1-C), respectively, as well as tiny grey and whitish colonies displaying the SCV phenotype (Figure 1-D). Figure 1-D also shows the hyper-splitting phenomenon of the SCV phenotype into the colony morphotypes shown in figures 1-A–C.

Initially, largely antibiotic-susceptible *K. pneumoniae* phenotypes exhibiting whitish and grey colony morphologies on Columbia agar plates were isolated from the urine sample. Following antibiotic treatment, MDR *K. pneumoniae* strains displaying the normal colony size were isolated from tissue samples, again characterized by subsequent whitish or grey colony formations. Subsequently, SCVs of *K. pneumoniae* were isolated from tissue samples. Subcultivation of different SCV colonies consistently yielded a division into four distinct colony morphotypes including one SCV phenotype that resembled the initial SCV, along with three normal-sized phenotypes distinguished by variations in colony color and visual attributes. While normal-sized phenotypes exhibited stability following each round of re-cultivation, SCV isolates displayed instability and recurrently diverged into the four phenotypes described above. We have designated the emergence of these multiple phenotypes as “hyper-splitting”. Despite minor variations in MIC values, these “hyper-splitting” phenotypes exhibited multidrug resistance (Table 1).

Except for isolates 1-A and 1-B, all isolates were resistant to the tested carbapenems. Initially, during routine diagnosis, isolate 2-B was found to be carbapenem-resistant by VITEK^®^ 2 AST, and to harbor *bla*_OXA-48_ gene by LAMP. After subcultivation of this isolate for MIC determination, this resistance disappeared and the isolate became susceptible to all tested beta-lactam antibiotics except piperacillin. Only isolates 1-A and 1-B were susceptible to piperacillin, and only isolate 4-B was not resistant to the cephalosporins tested. Interestingly, only isolates 4-A and 5-A, which demonstrated the SCV phenotype, were resistant to amikacin and trimethoprim-sulfamethoxazole. Another remarkable finding was the observed increase in the MIC values of cefiderocol and trimethoprim-sulfamethoxazole over time (Table 1).

Whole-genome sequence (WGS) analysis revealed that all isolates belonged to sequence type (ST) 147. Lipopolysaccharide antigen (O) loci were O1/O2v1 and capsule biosynthesis (KL) loci were KL64 for all isolates except isolate 4-D, which could not be assigned, as it missed most genes of this locus. Isolates 1-A, 1-B and 2-B showed lower Kleborate resistance score than the other isolates (resistance: 0 vs. 2). The resistance score of 0 indicates that the isolate(s) did not carry any genes for extended-spectrum beta-lactamases (ESBL) or carbapenemases and a score of 2 correlated with the presence of carbapenemase genes without colistin resistance genes (35). In accordance with the resistance scores, we detected several beta-lactamase genes, such as *bla*_SHV-11_, *bla*_TEM-1_ and *bla*_OXA-9_, ESBL genes, such as *bla*_CTX-M-15_ and *bla*_OXA-1_, and the carbapenemase genes *bla*_NDM-1_ and *bla*_OXA-48_. *bla*_SHV-11_ was found in all isolates whereas *bla*_TEM- 1_ and *bla*_OXA-9_ were present in all isolates except 1-A and 1-B. However, *bla*_CTX-M-15_ was not found in isolate 4-A. Genes associated with sulphonamide (*sul1*) and chloramphenicol (*catB3*) resistance were also detected in all isolates except 1-A, 1-B and 2-B (Table S1). Note that we did not detect any common cefiderocol resistance genes.

The isolates exhibited clonality as emphasized by the low number of SNPs among them (Table S1). Especially isolates from the same time point showed no difference in the core genome alignment (5,360,988 bp) with the exception of 2-A and 2-B (six SNPs) and 5-D (one additional SNP compared to 5-A–C). The largest distance with 17 SNPs was between 2-A and 5-D (Table S1).

## Discussion

When evaluating the results, we can roughly identify three distinct outcomes. The first significant observation concerns the emergence of resistance development chronologically within a *K. pneumoniae* strain, originating from a patient subjected to continuous, uninterrupted antibiotic intervention. This scenario promptly elicits contemplation of the subject concerning within-host adaptive evolution of bacteria. In fact, in-host resistance evolution, either due to plasmid mediation or chromosome mutations, has been observed even shortly after the initiation of antimicrobial treatment (38).

The second notable observation in our study is the occurrence of SCVs from patient specimens following the detection of normal-sized morphotypes. SCVs demonstrate remarkable abilities to invade and persist within host cells, thus evading the surveillance mechanisms of the immune system (39). The existence of SCVs, mostly observed in *Staphylococcus* spp., has been documented since the onset of the 20th century and has gained increasing attention due to its potential implications for both clinical and basic research (12, 40). Regarding the SCVs of Gram-negative bacteria, studies have particularly focused on *Burkholderia* and *Pseudomonas* spp. isolated from CF patients (18, 41, 42). However, there are only sparse data on the occurrence of SCV in *Klebsiella* spp. (23-27).

Basically, SCVs have been determined as a subpopulation characterized by their distinct phenotypic properties, such as atypical colony morphologies including the reduced colony size (43). Their decreased growth rate is thought to contribute to their inherent resistance, given that the decelerated growth dynamics potentially hinder the effectiveness of antibiotics geared towards rapidly proliferating cell populations (44). Furthermore, this phenomenon concurrently signifies decreased metabolic activity, which may engender modifications in cell wall permeability, drug uptake, or the modulation of efflux pump expression (45).

For electron transport chain-defective staphylococcal SCVs, lower efficacy of aminoglycosides known to be taken up through electrical potential across the cytoplasmic membrane (`ΔΨ)` was demonstrated, which is attributable to low `ΔΨ` (46). These alterations could collectively contribute to enhancing resistance patterns. In this study, we observed an increase in the MIC values of amikacin, cefiderocol, and trimethoprim-sulfamethoxazole in the isolates recovered over time. This MIC increase was especially pronounced for amikacin in SCV phenotypes. Moreover, most antibiotics penetrate into host cells poorly, so the concentrations required to kill intracellularly persistent SCVs cannot be achieved (12).

SCVs, known for their inducible formation through *in vitro* processes involving various agents, including antibiotics (23), have exhibited a propensity for increased persistence and adaptability when confronted with challenging environments (47). An enhanced ability to form biofilms on biotic and abiotic surfaces has been shown for SCVs of different bacterial species (41, 48-51). The substantial implication of SCVs extends to their involvement in biofilm development, as biofilms effectively shield bacteria from harsh host environments, thereby complicating the elucidation of drug resistance mechanisms within biofilm structures (52). Biofilms not only confer protection against host immune defenses but also serve as reservoirs for persistent infections and recurrent episodes (53). The impact of SCV phenotype on biofilm formation in in *Klebsiella* remains to be elucidated in further studies.

Furthermore, the emergence of SCVs could plausibly be due to selection pressure from antibiotic regimens or other host-associated factors, e.g., host cationic peptides. Consistent with the case that was the subject of our study, the higher frequency of SCVs in isolates from chronic and recurrent infections compared to acute infections suggests a potential role for these variants in evading host immune responses and antimicrobial treatments (12). In the context of our study, the emergence of SCVs after the initiation of cefiderocol treatment while already undergoing antibiotic therapy could be construed as a form of *in vivo* or *in host* induction.

The third noteworthy finding from our study underscores the inherent instability of SCVs. This dynamic interplay between stable and unstable SCVs is still poorly understood and its elucidation may contribute to a deeper understanding of their role in infection in general and persistence phenomena in particular (54). Despite comprehensive explorations largely focusing on staphylococci, a lack of investigations concerning *Klebsiella* spp. persists, and requires attention.

The observed instability among SCVs, combined with distinct antibiotic susceptibility profiles across phenotypes, increases the significance of investigating SCV plasticity (43). Stable SCVs represent a long-term adaptation strategy, whereas their unstable counterparts may arise as stress-induced variants that result from rapid adaptation to fluctuating environments (14, 55, 56). This inherent instability potentially serves as a mechanism for evading host immune responses and circumventing antibiotic interventions (55). Furthermore, the involvement of epigenetic modifications, including alterations in DNA methylation patterns, could significantly influence SCV stability (57). In addition, regulatory systems, such as two-component systems and quorum sensing, play a crucial role in SCV formation by modulating bacterial behavior and adaptation. Disruption or dysregulation of these systems could lead to the emergence of SCVs with altered phenotypic properties (58). Due to instability, slow-growing SCVs may generate mutants that exhibit a faster growth rate than usual (59). In instances of reversion to the wild type, rapidly growing mutant revertants may demonstrate either the loss or preservation of antibiotic resistance (59).

A high mutation rate might favor the emergence of SCVs (20) and also explain the emergence of antibiotic resistance as a result of antibiotic selective pressure and the adaptation of hypermutable strains in patients, especially CF patients (19). CF-like chronic infections have been shown to specifically contribute to the development of bacterial mutations (60). Hypermutation could result in a subpopulation of bacteria that temporarily does not grow, thus leading to persistence (61). Additionally, an increase in the prevalence of mutator bacterial strains with deficient DNA mismatch repair (MMR) system has been detected in CF patients, who are used as a reservoir for mutation (62). To our best knowledge, we were unable to identify any instance in the available literature wherein a solitary SCV colony has given rise to four distinct colonies exhibiting disparate morphologies. Accordingly, we suggest the designation “phenotypic hyper-splitting” for this distinctive phenomenon.

We described in this study unprecedented phenotypic attributes and primarily focused on *in vitro* experiments. Therefore, the clinical relevance of our findings necessitates validation through animal models and clinical sample analyses. In this context, macrophage and neutrophil assays would be valuable for assessing both the extent of immune response and the presence of persistent cells. Moreover, the determination of the auxotrophism (13, 17) of *K. pneumoniae* SCVs and of the molecular mechanisms that drive SCV formation and the resulting antibiotic resistance in this species require further investigation. Integrating a comprehensive range of approaches encompassing genomics, transcriptomics, and proteomics, the utilization of experimental evolutionary models can yield valuable insights into the genetic determinants and regulatory networks orchestrating SCV phenotypes.

The genomic analysis conducted in this study has revealed clonality among all 14 isolates. Further exploration is warranted to uncover the intricate molecular mechanisms underlying phenotypic hyper-splitting and to elucidate the potential pathogenic implications of this phenomenon. To better understand the formation of the SCV phenotype especially in Gram-negative pathogens, efforts need to be intensified (i) to improve the detection and characterization of SCVs recovered from clinical samples and (ii) to elucidate their clinical impact.

## Data availability

The data for this study have been deposited in the European Nucleotide Archive (ENA) at EMBL-EBI under accession number PRJEB71325 (https://www.ebi.ac.uk/ena/browser/view/PRJEB71325).

## Supplemental material

**Table S1**. Core SNP distance matrix. The complete core genome alignment (gaps and ambiguous bases removed) contained 5,360,988 bp. The reference sequence for alignment was 1-A.

## Acknowledgments

We are grateful to Betty Nedow, Katrin Darm, Maysem Al-Baldawi and Sara-Lucia Wawrzyniak for their technical assistance.

## Funding

This study was funded in part by a grant from the European Regional Development Fund (ERDF) to K.B. (grant number GHS-20-0010). Support was also obtained from a grant from the Federal Ministry of Education and Research (BMBF, Germany) to KSc entitled “Disarming pathogens as a different strategy to fight antimicrobial-resistant Gram-negatives” (01KI2015).

## Conflict of interest

The authors declare no conflict of interest.

